# CRISPR-Cas9 generated Pompe knock-in murine model exhibits early-onset hypertrophic cardiomyopathy and skeletal muscle weakness

**DOI:** 10.1101/780320

**Authors:** Jeffrey Y. Huang, Shih-Hsin Kan, Emilie K. Sandfeld, Nancy D. Dalton, Anthony D. Rangel, Yunghang Chan, Jeremy Davis-Turak, Jon Neumann, Raymond Y. Wang

## Abstract

Infantile-onset Pompe Disease (IOPD), caused by mutations in lysosomal acid alpha-glucosidase (*Gaa*), manifests rapidly progressive fatal cardiac and skeletal myopathy incompletely attenuated by synthetic GAA intravenous infusions. The currently available murine model does not fully simulate human IOPD, displaying skeletal myopathy with late-onset hypertrophic cardiomyopathy. Bearing a Cre-*LoxP* induced exonic disruption of the murine *Gaa* gene, this model is also not amenable to genome-editing based therapeutic approaches. We report the early onset of severe hypertrophic cardiomyopathy in a novel murine IOPD model generated utilizing CRISPR-Cas9 homology-directed recombination to harbor the orthologous *Gaa* mutation c.1826dupA (p.Y609*), which causes human IOPD. We demonstrate the dual sgRNA approach with a single-stranded oligonucleotide donor is highly specific for the *Gaa*^c.1826^ locus without genomic off-target effects or rearrangements. Cardiac and skeletal muscle were deficient in *Gaa* mRNA and enzymatic activity and accumulated high levels of glycogen. The mice demonstrated skeletal muscle weakness but did not experience early mortality. Altogether, these results demonstrate that the CRISPR-Cas9 generated *Gaa*^c.1826dupA^ murine model recapitulates hypertrophic cardiomyopathy and skeletal muscle weakness of human IOPD, indicating its utility for evaluation of novel therapeutics.

## Introduction

Generation of transgenic murine knock-in models of human disease once relied solely upon targeted insertion of the desired sequence via Cre-*Lox* recombination and embryonic stem cell implantation. Though this strategy has resulted in many successful murine model systems, it is labor-intensive, time-consuming, and expensive. The advent of genome editing via engineered nucleases, especially clustered regularly interspaced short palindromic repeats (CRISPR)-based systems has allowed for a potentially accurate, efficient, and relatively inexpensive alternative to the traditional method of transgenic knock-in model generation^1,2^.

As an intriguing example of disorders that may uniquely benefit from genome editing, inherited metabolic disorders (IMDs) are a diverse group of genetic diseases affecting the proper breakdown or synthesis of essential compounds such as carbohydrates, amino acids, or organic acids. Many of these disorders are caused by single gene defects that alter the expression and/or activity of critical metabolic enzymes. Given the monogenic nature of IMD pathogenesis, this class of genetic disorders is quickly becoming an area of high interest for CRISPR-mediated genome editing therapeutics^3,4,5^.

Pompe disease, caused by the deficiency of acid α-glucosidase (GAA; EC 3.2.1.20), is characterized by lysosomal accumulation of glycogen in body tissues, primarily cardiac and skeletal muscle. Muscle lysosomal glycogen storage results in muscle weakness varying in age of onset and severity according to residual GAA enzymatic activity. Infantile-onset Pompe disease (IOPD), caused by nearly absent GAA enzyme, typically manifests in the first two months of life with progressive and severe hypertrophic cardiomyopathy, heart failure, hypotonia, respiratory failure, and death within the first 14 months of life^6^.

Pompe disease can be treated with intravenous enzyme replacement therapy (ERT) using recombinant human acid α-glucosidase (rhGAA) enzyme, which significantly reduces cardiac hypertrophy and increases overall and ventilator-free survival^7^. Unable to endogenously synthesize GAA, Pompe patients are infused indefinitely and may produce an anti-rhGAA antibody response that may limit or neutralize treatment efficacy. Regardless of immune response, glycogen storage, autophagic buildup, and fibrosis within skeletal myocytes are observed even in early-treated IOPD patients^8,9^.

Consequently, a phenotype of sensorineural hearing loss, central nervous system white matter abnormalities, slowly progressive muscle weakness and delayed mortality is now observed in rhGAA-treated survivors with IOPD^10,11^.

The limitations of current Pompe disease treatment underscore the necessity of new therapeutic development. CRISPR-based therapeutic strategies may address the impermanence of ERT, effecting permanent, highly-specific somatic correction of genomic *Gaa* mutations within myocytes and subsequent intramuscular, endogenous synthesis of enzyme. Reduced GAA enzyme within the bloodstream may also mitigate the immunogenicity of intravenous ERT.

First, though, an animal model with molecular, biochemical, physiological and functional analogy to human Pompe disease must be developed. Currently, there are knockout murine models of Pompe disease featuring large exonic disruptions in the *Gaa* gene^12,13^. The most widely used preclinical model of Pompe disease is the Raben knockout mouse (B6;129-*Gaa*^tm1Rabn/J^) that demonstrates survival into adulthood with muscle glycogen storage and progressive muscle weakness^13^. This model bears a neomycin resistance cassette (~800bp) in exon 6 of the *Gaa* gene, not a homolog of a human *Gaa* mutation, complicating preliminary efforts at *in vivo* genome correction. Here, we report the successful generation of a *Gaa*^c.1826dupA^ (p.Y609*)^14^ murine knock-in model of Pompe disease utilizing a novel, dual-single guide RNA approach flanking the intended *Gaa* insertion site, and early characterization that demonstrates GAA enzymatic deficiency, hypertrophic cardiomyopathy, muscular glycogen storage and pathology recapitulating human Pompe disease.

## Results

### Dual overlapping gRNA approach achieves highest HDR levels in vitro

Three guide RNAs (*Gaa*^c.1826^ gRNA-1,2,3) were selected based on best predicted on-target and off-target scoring and proximity of expected cut site to *Gaa*^c.1826^ target locus (**Figure 1A**). To determine *in vitro* on-target editing activity and homology-directed repair (HDR) efficiency, *Gaa*^c.1826^ gRNA expression vectors and respective single-stranded donor oligonucleotides (ssODN) were nucleofected into C2C12 mouse myoblasts. On-target editing activity and HDR efficiency was highly dependent on target sequence (**Table 1**). For single gRNA approach, *Gaa*^c.1826^ gRNA-3 demonstrated the highest on-target editing activity (47.9±0.1%) with *Gaa*^c.1826^ gRNA-2 achieving the best HDR efficiency (7.7±1.4%). Given that multiple gRNAs with overlapping sequences are known to enhance CRISPR/Cas9-mediated knock-in efficiency^15^, we evaluated whether a dual overlapping gRNA approach could improve our on-target editing activity and HDR efficiency. We chose to test *Gaa*^c.1826^ gRNAs-2 and −3 at an equimolar ratio as they achieved the highest levels of on-target editing activity and HDR efficiency. Furthermore, *Gaa*^c.1826^ gRNAs-2 and −3 are a sense-antisense pair with fully overlapping target sequences thereby reducing the likelihood of added off-target activity. We found that the dual overlapping gRNA approach achieved high on-target editing activity (41.0±4.4%) with the highest overall HDR efficiency (12.6±2.9%). Further testing will need to be performed to confirm that this dual overlapping gRNA approach can be broadly applied to increase HDR efficiency in other target loci and cell lines.

**Figure 1:**
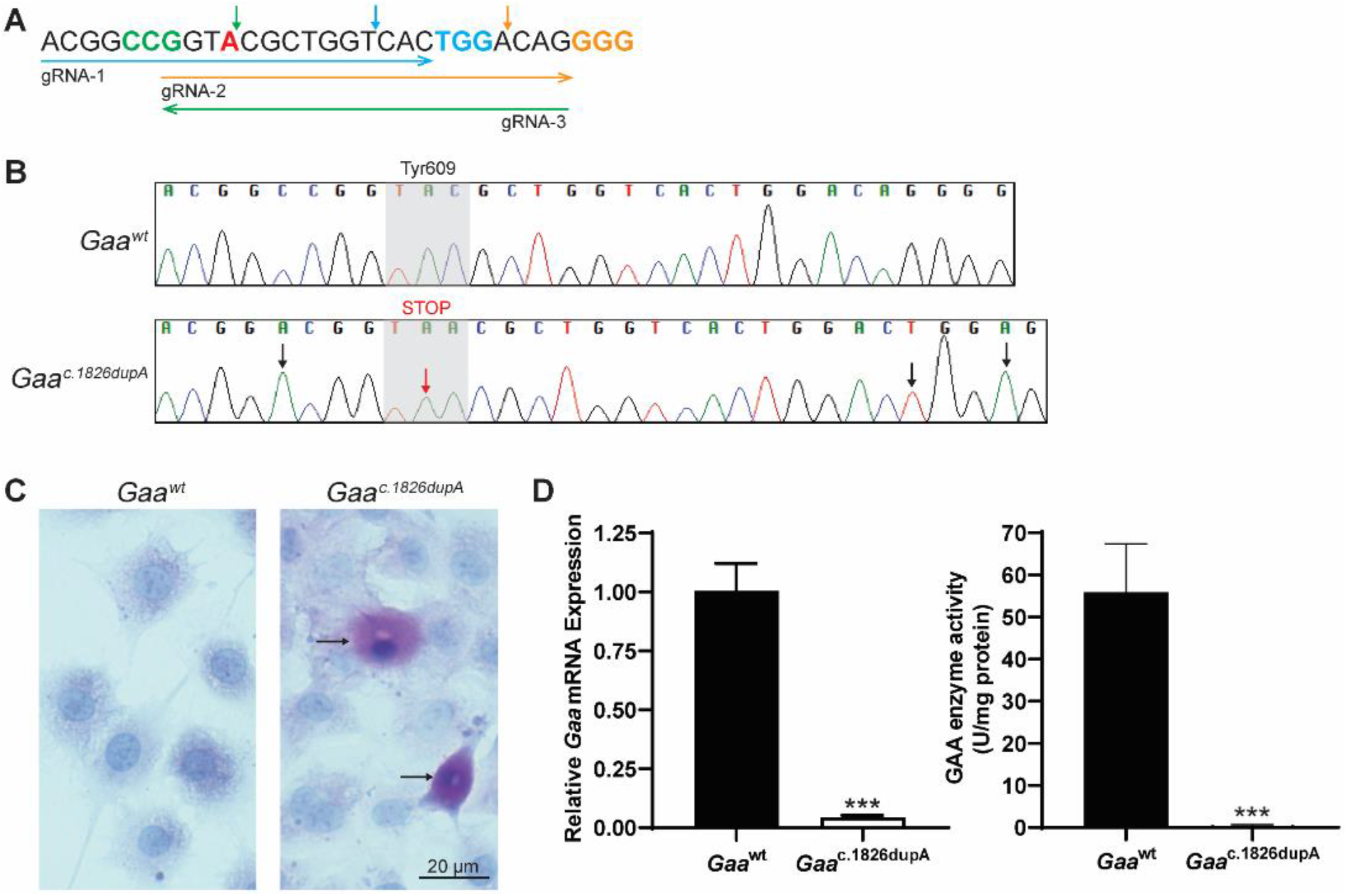
Generation of *Gaa*^c.1826dupA^ C2C12 clonal cell line. **(A)** Sequences of guide RNAs targeting *Gaa*^c.1826^ target locus. Arrow direction indicates directionality of guide RNA: sense (right) or antisense (left). gRNA-2 and gRNA-3 are reverse complements and were selected for dual overlapping gRNA strategy. Protospacer adjacent motifs (PAM; NGG) are highlighted in color corresponding to guide RNA arrow. *Gaa*^c.1826^ locus for targeted insertion of additional adenine nucleotide is highlighted in red. Expected Cas9 nuclease cut sites are shown as vertical arrows in color corresponding to guide RNA arrow. **(B)** Sanger sequencing chromatograms of control (*Gaa*^wt^) and clonal knock-in (*Gaa*^c.1826dupA^) C2C12 myoblast genomic DNA at *Gaa*^c.1826^ locus. Black arrows indicate silent mutations at PAM sites (*Gaa*^c.1821C>A^, *Gaa*^c.1845G>A^) or gRNA seed region (*Gaa*^c.1842A>T^). Red arrow indicates desired knock-in mutation (*Gaa*^c.1826dupA^). Gray shaded region indicates amino acid change at position 609 from tyrosine (TAC) to stop codon (TAA). **(C)** Periodic-acid Schiff (PAS) staining of control (*Gaa*^wt^) and clonal knock-in (*Gaa*^c.1826dupA^) C2C12 myoblasts. Fixed cells were PAS stained (purple-magenta) and hematoxylin counterstained (blue). Only *Gaa*^c.1826dupA^ knock-in myoblasts display accumulation of PAS staining (see arrows). Representative images were captured on a bright-field microscope (Olympus) at 40x objective magnification. **(D)** (Left panel) *Gaa* mRNA expression in *Gaa*^wt^ and *Gaa*^c.1826dupA^ C2C12 myoblasts. *Gaa*^c.1826dupA^ myoblasts display markedly reduced *Gaa* transcript levels relative to *Gaa*^wt^ myoblasts. Relative *Gaa* expression levels were measured by TaqMan probe-based quantitative real-time PCR using comparative C_t_ method of target gene (*Gaa*) to reference gene (*Gapdh*). Data are generated from three independent experiments and comparisons were analyzed with unpaired two-tailed t-test. ***p<0.001. (Right panel) GAA enzymatic activity in *Gaa*^wt^ and *Gaa*^c.1826dupA^ C2C12 myoblasts. *Gaa*^c.1826dupA^ myoblasts display markedly reduced GAA activity levels relative to *Gaa*^wt^ myoblasts. GAA enzymatic activity (fluorescent units) was measured using a fluorometric 4-MU α-D-glucoside assay and normalized to amount of sample protein. Data are generated from three independent experiments and comparisons were analyzed with unpaired two-tailed t-test. ***p<0.001.

### Generation & characterization of Gaa^c.1826dupA^ knock-in C2C12 cell line

We used the dual overlapping *Gaa*^c.1826^ gRNA strategy (**Figure 2A,B**) followed by puromycin-resistant selection to isolate *Gaa*^c.1826dupA^ knock-in C2C12 clonal cells. Sanger sequencing results confirmed presence of desired *Gaa*^c.1826dupA^ knock-in mutation as well as silent protospacer adjacent motif (PAM) and seed region mutations to prevent gRNA editing of the donor template (**Figure 1B**). *Gaa*^c.1826dupA^ knock-in cells exhibited enhanced Periodic acid-Schiff (PAS) staining – a marker of glycogen accumulation – relative to *Gaa*^wt^ cells (**Figure 1C**). *Gaa*^c.1826dupA^ knock-in cells display a 96% reduction in *Gaa* transcript levels and GAA enzymatic activity was completely abolished (**Figure 1D)**. Together, these results demonstrate that the dual overlapping gRNA approach can increase overall HDR efficiency *in vitro* thus improving the probability of isolating clonal cells with a desired knock-in mutation. Moreover, our *Gaa*^c.1826dupA^ knock-in cell line exhibits molecular and biochemical analogy to human Pompe disease thereby validating its use as an *in vitro* model.

**Figure 2:**
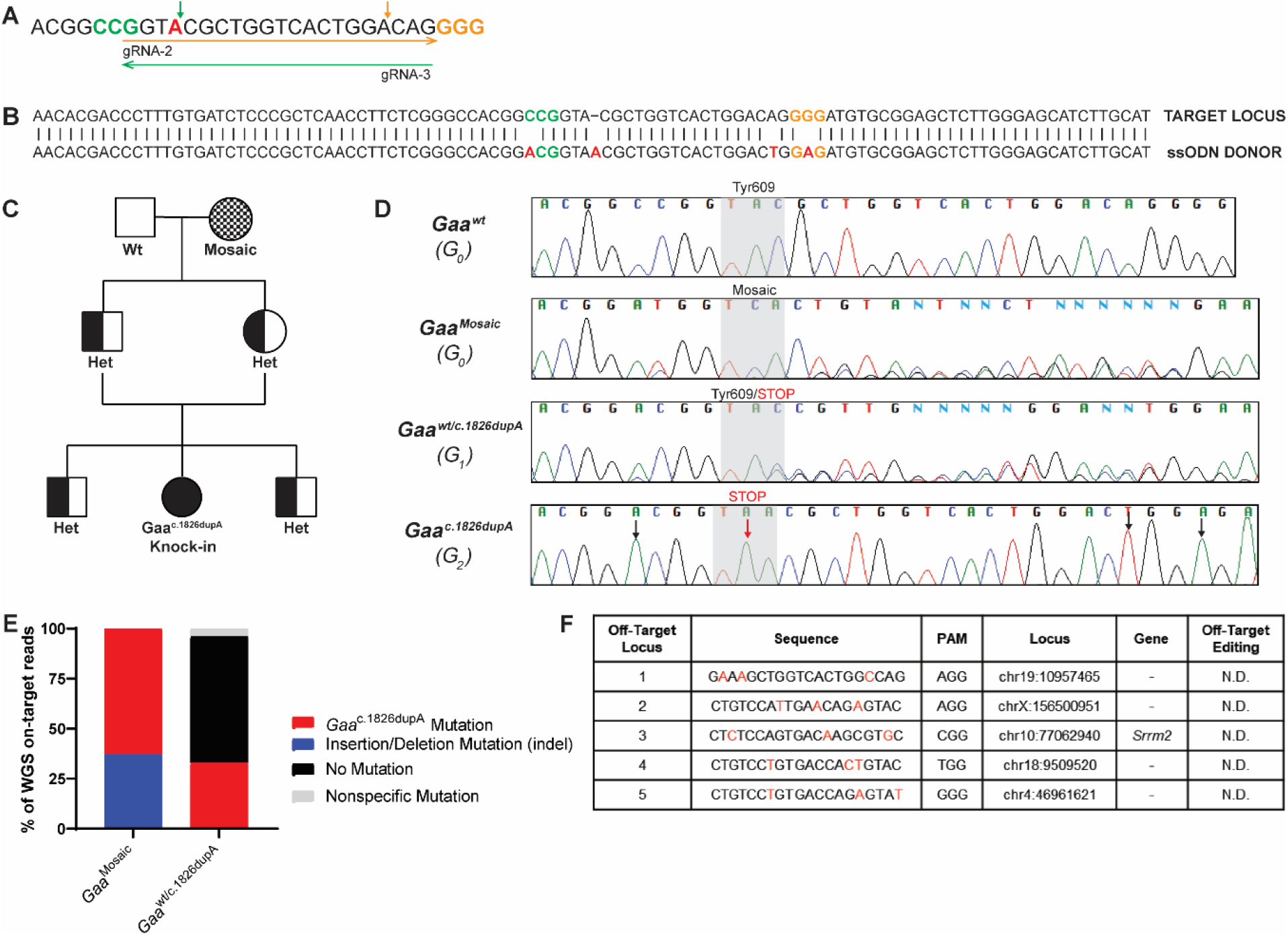
Generation of *Gaa*^c.1826dupA^ knock-in transgenic mouse line. **(A)** Dual overlapping guide RNA approach targeting *Gaa*^c.1826^ target locus. Arrow direction indicates whether guide RNA is sense (right) or antisense (left). Protospacer adjacent motifs (PAM; NGG) are highlighted in color corresponding to guide RNA arrow. *Gaa*^c.1826^ locus for targeted insertion of additional adenine nucleotide is highlighted in red. Expected Cas9 nuclease cut sites are shown as vertical arrows in color corresponding to guide RNA arrow. **(B)** Sequence of single-stranded donor oligonucleotide (ssODN) for targeted integration of *Gaa*^c.1826dupA^ knock-in mutation. PAM motifs are indicated in either gold (gRNA-2) or green (gRNA-3). Silent mutations at PAM sites (*Gaa*^c.1821C>A^, *Gaa*^c.1845G>A^), gRNA seed region (*Gaa*^c.1842A>T^) and desired knock-in mutation (*Gaa*^c.1826dupA^) are highlighted in red. **(C)** Pedigree diagram of mating scheme to isolate *Gaa*^c.1826dupA^ knock-in allele from mosaic CRISPR-generated founder mouse for generation of homozygous *Gaa*^c.1826dupA^ knock-in mice. Males are represented as squares and females are represented as circles. **(D)** Sequencing chromatograms of control (*Gaa*^wt^), founder (*Gaa*^Mosaic^), heterozygous (*Gaa*^wt/c.1826dupA^) and homozygous knock-in (*Gaa*^c.1826dupA^) mice. Black arrows indicate silent mutations at PAM sites (*Gaa*^c.1821C>A^, *Gaa*^c.1845G>A^) or gRNA seed region (*Gaa*^c.1842A>T^). Red arrow indicates desired knock-in mutation (*Gaa*^c.1826dupA^). Gray shaded region indicates amino acid(s) at position 609 for each mouse. **(E)** WGS on-target analysis (>50x read depth) of *Gaa*^c.1826^ locus in G_0_ founder (*Gaa*^Mosaic^) and G_1_ heterozygous (*Gaa*^wt/c.1826dupA^) mice. WGS analysis demonstrates highly efficient on-target genome-editing in *Gaa*^Mosaic^ mice and successful passing on of desired Gaa^c.1826dupA^ mutation to G_1_ heterozygous mouse. Stacked bar graphs indicate % of on-target reads for each mutation event or non-event. **(F)** List of the 5 most similar off-target sequences to *Gaa*^c.1826^ gRNA-2 & 3 used for WGS off-target screening. Nucleotide letters shown in red are the individual mismatches between off-target and *Gaa*^c.1826^ gRNA target sequences. (−) indicates off-target sequence is found on intronic region. N.D. = none detected.

### Generation & characterization of Gaa^c.1826dupA^ transgenic mice

We next applied the dual overlapping *Gaa*^c.1826^ gRNA strategy *in vivo* (**Figure 2A,B**) to generate transgenic *Gaa*^c.1826dupA^ knock-in mice via pronuclear injection of C57BL/6NJ single-cell embryos by standard methods^16^. The dual overlapping *Gaa*^c.1826^ gRNA strategy achieved 66.7% on-target editing activity (founder mice positive for any *Gaa* mutation) and 25% HDR efficiency (founder mice positive for *Gaa* ^c.1826dupA^ mutation) (**Table 2**). Following founder mice genotyping, we selected a founder with the lowest levels of mosaicism - as determined by TIDE^17^ and TIDER^18^ analysis – for mating and segregation of *Gaa* ^c.1826dupA^ mutation. We successfully generated a homozygous *Gaa* ^c.1826dupA^ knock-in mouse after 2 generations of breeding (**Figure 2C**). Sequencing results confirmed presence of desired *Gaa*^c.1826dupA^ knock-in mutation as well as silent PAM and seed region mutations in G_0_ founder (*Gaa*^Mosaic^), G_1_ heterozygous (*Gaa*^wt/c.1826dupA^) and G_2_ homozygous knock-in (*Gaa*^c.1826dupA^) mice (**Figure 2D**). To determine the extent of genomic mosaicism in our transgenic mice, we performed whole genome sequencing at >50x coverage of the G_0_ founder (*Gaa*^Mosaic^), G_1_ heterozygous (*Gaa*^wt/c.1826dupA^) and G_0_ wild-type (*Gaa*^wt^) mice. On-target locus alignment of *Gaa*^Mosaic^ to *Gaa*^wt^ demonstrates 63% *Gaa*^c.1826dupA^ knock-in mutation (32/51 reads) and 37% indels (19/51 reads). Alignment of *Gaa*^wt/c.1826dupA^ to *Gaa*^wt^ resulted in 33% *Gaa*^c.1826dupA^ knock-in mutation (17/51 reads), 63% no mutation (32/51 reads) and 4% nonspecific mutation (2/51 reads) (**Figure 2E**). Together, these results demonstrate that the desired *Gaa*^c.1826dupA^ knock-in mutation efficiently integrated into our founder mouse genome and was successfully passed onto its progeny.

To screen for off-target integration of the donor template, we searched for called single nucleotide variants (SNVs) that had the unique donor template motif. We then repeated this step for the reverse complement and found that the only positive result was the intended mutation at the *Gaa*^c.1826^ target locus. Next, we screened the top 5 genomic regions predicted by GT-Scan^19^ to be potential off-target sites (**Figure 2F**). There were no detected SNVs within 500bp of these sites.

Altogether, these results suggest that the dual overlapping gRNA approach is an efficient strategy to generate transgenic *Gaa*^c.1826dupA^ knock-in mice and did not result in any detectable off-target activity in the genomes of our founder mouse and its progeny.

### Gaa transcript levels, GAA enzyme activity and glycogen load in Gaa^c.1826dupA^ transgenic mice

Knock-in of the *Gaa*^c.1826dupA^ mutation leads to a frameshift in the *Gaa* coding sequence resulting in a premature stop codon at amino acid position 609 (p.Tyr609*). Nonsense mediated decay of transcripts containing premature stop codons prevents expression of truncated and potentially deleterious proteins. Thus, we assessed nonsense mediated decay by measuring *Gaa* transcript levels from *Gaa*^*c.1826dupA*^ transgenic and wild-type mouse tail samples using quantitative real-time PCR (qRT-PCR). Transgenic mice demonstrated a gene dose-dependent decrease in *Gaa* transcript levels relative to *Gaa*^*wt*^ (WT) mice (**Figure 3A**). *Gaa*^*c.1826dupA*^ homozygous (KI) and *Gaa*^wt/c.1826dupA^ heterozygous (HET) mice exhibited a 75% and 46% reduction in *Gaa* transcript levels relative to WT, respectively. To compare these results with the commercially available *Gaa*^tm1Rabn^ knockout (KO) mice, we found an 81% reduction in KO *Gaa* transcript levels relative to WT. Notably, there was no significant difference in *Gaa* transcript levels between KI and KO mice.

**Figure 3:**
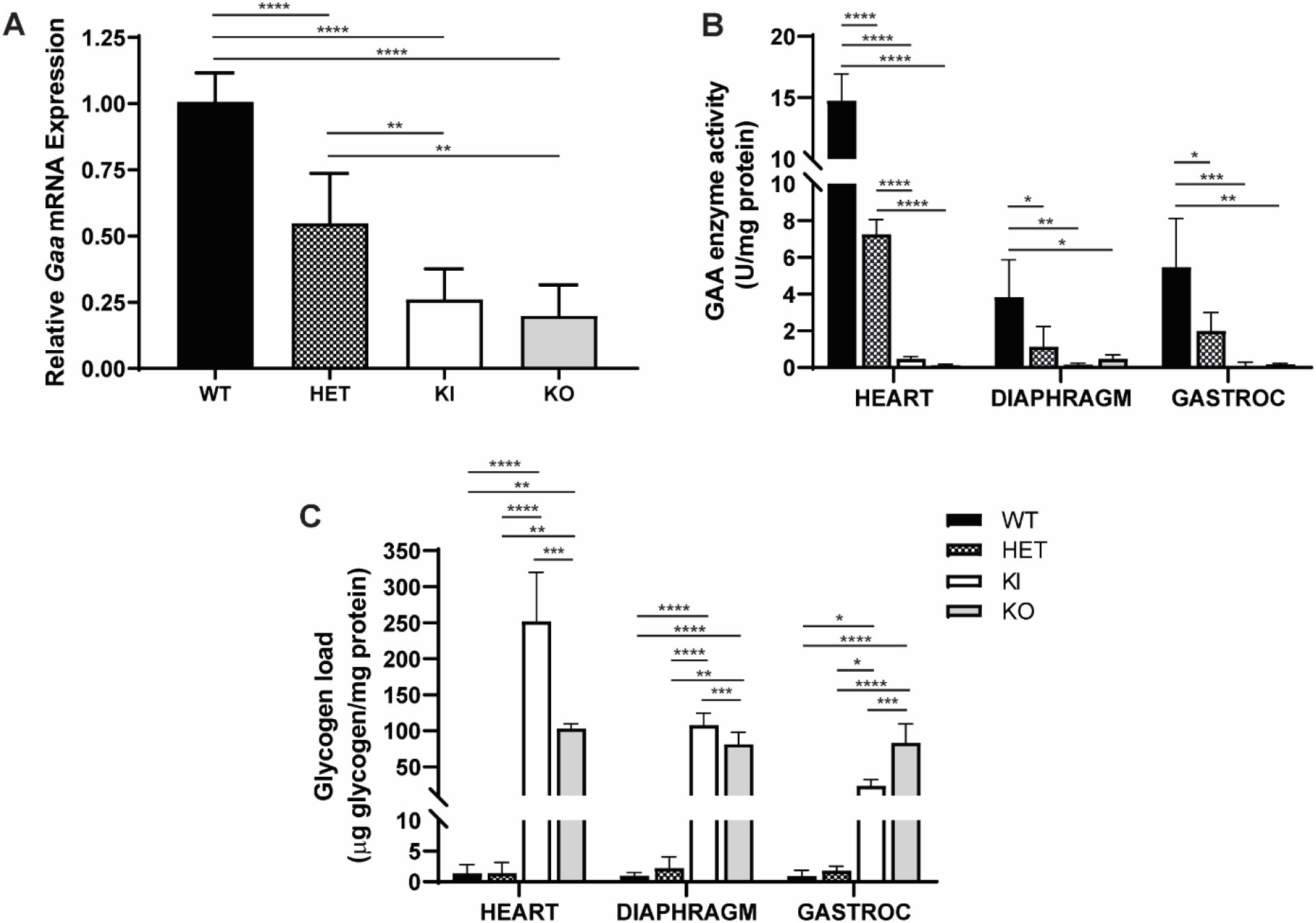
*Gaa*^c.1826dupA^ knock-in mice show reduced GAA expression and enzymatic activity. **(A)** *Gaa* mRNA expression in tail biopsies of 3-week old *Gaa*^wt^ (WT, n=8), *Gaa*^wt/c.1826dupA^ (HET, n=12), *Gaa*^c.1826dupA^ (KI, n=5) and *Gaa*^tm1Rabn^ (KO, n=3) mice. KI and KO mice display significantly reduced *Gaa* transcript levels relative to WT and HET mice. Relative *Gaa* expression levels were measured by TaqMan probe-based quantitative real-time PCR using comparative C_t_ method of target gene (*Gaa*) to reference gene (*Gapdh*). Data are generated from two independent experiments and all comparisons were analyzed using one-way ANOVA with Tukey post-hoc test. **p<0.01, ****p<0.0001. **(B)** GAA enzymatic activity in the heart, diaphragm and gastrocnemius muscle of *Gaa*^wt^ (WT, n=5), *Gaa*^wt/c.1826dupA^ (HET, n=5), *Gaa*^c.1826dupA^ (KI, n=5) and *Gaa*^tm1Rabn^ (KO, n=3) mice. KI and KO mice display significantly reduced GAA enzyme activity levels in heart (relative to WT and HET) as well as diaphragm and gastrocnemius muscle (relative to WT). GAA enzymatic activity (fluorescent units) was measured using a fluorometric 4-MU α-D-glucoside assay and normalized to amount of sample protein. Data are generated from three independent experiments and all comparisons were analyzed using one-way ANOVA with Tukey post-hoc test. ^*^p<0.05, **p<0.01, ***p<0.001, ****p<0.0001. **(C)** Glycogen levels were measured in the heart, diaphragm and gastrocnemius muscle of *Gaa*^wt^ (WT, n=5), *Gaa*^wt/c.1826dupA^ (HET, n=5), *Gaa*^c.1826dupA^ (KI, n=5) and *Gaa*^tm1Rabn^ (KO, n=3) mice using a colorimetric assay. KI and KO mice display significantly elevated glycogen levels relative to WT and HET in heart, diaphragm and gastrocnemius muscle. Glycogen amount is normalized to amount of sample protein. Data are generated from three independent experiments and all comparisons were analyzed using one-way ANOVA with Tukey post-hoc test. ^*^p<0.05, **p<0.01, ***p<0.001, ****p<0.0001.

Next, we measured GAA enzyme activity levels in the heart, diaphragm and gastrocnemius muscle of WT, HET, KI and KO mice. For all tissues, transgenic mice demonstrated a gene dose-dependent decrease in GAA enzyme activity levels relative to WT mice (**Figure 3B**). Heart GAA activity was reduced by 97% and 51% reduction in KI and HET mice relative to WT, respectively. Diaphragm GAA activity was reduced by 96% and 71% in KI and HET mice relative to WT, respectively. Gastrocnemius muscle GAA activity was reduced by 98% and 64% in KI and HET mice relative to WT, respectively. We also assessed GAA activity in KO mice and found a 99%, 88% and 97% reduction in heart, diaphragm and gastrocnemius GAA activity relative to WT, respectively. Notably, there was no significant difference in GAA enzymatic activity between KI and KO mice for all tissue types.

We then measured glycogen load in the heart, diaphragm and gastrocnemius muscle of WT, HET, KI and KO mice. For all tissues, KI mice demonstrated a substantial increase in glycogen load relative to WT mice (**Figure 3C**). Glycogen levels were increased 185-fold, 108-fold and 28-fold in the heart, diaphragm and gastrocnemius of KI mice relative to WT, respectively. We also measured glycogen load in KO mice and found a 76-fold, 82-fold and 89-fold increase in heart, diaphragm and gastrocnemius glycogen levels relative to WT, respectively. Interestingly when compared to KI, KO glycogen load was 2.4-fold lower, 1.3-fold lower and 3.2-fold higher in heart, diaphragm and gastrocnemius, respectively.

Taken together, these results demonstrate that our *Gaa*^c.1826dupA^ knock-in mouse model exhibits molecular and biochemical analogy to the established preclinical model of Pompe disease (*Gaa*^tm1Rabn^) and is an appropriate *in vivo* model for genome-based therapeutic evaluation.

### Autophagy in Gaa^c.1826dupA^ transgenic mice

Progression of Pompe disease pathology in affected tissue is known to involve dysfunctional autophagy, resulting in the accumulation of autophagic markers such as microtubule-associated protein light chain 3B (LC3B)^20^. To assess autophagy in *Gaa*^c.1826dupA^ transgenic mice, we performed Western blot analysis of LC3B protein expression in the heart, diaphragm and gastrocnemius muscle of 3-month old WT, HET, KI and KO mice. It is important to note that endogenous LC3B protein expression is detected as two bands by Western blot: 1) cytosolic LC3B-I (16kD) and 2) autophagosomal LC3B-II (14kD) which is conjugated with phosphatidylethanolamine (**Supp. Figure 1**). LC3B-I expression was increased 2.3-fold and 3.6-fold in the heart and gastrocnemius muscle of KI mice relative to WT (**Figure 4A**). LC3B-II expression was increased 3.2-fold in the heart of KI mice relative to WT and increased 4.8-fold in the gastrocnemius muscle of both KI and KO mice relative to WT (**Figure 4B**). In contrast, LC3B-I and LC3B-II expression levels in the diaphragm of KI or KO mice were not significantly altered relative to WT or HET mice. Given that LC3B-II expression is closely correlated with autophagosome formation, the ratio of LC3B-II to total LC3B expression is often used to monitor autophagy. Thus, we measured LC3B-II to total LC3B ratios in our samples and found a significant reduction in the diaphragm of KO mice relative to WT and KI mice (**Figure 4C**). Notably, LC3B-II to total LC3B ratios in the gastrocnemius muscle of KI and KO mice – which exhibited the highest fold increases in LC3B-II expression relative to WT – were 1.2-fold and 1.7-fold higher relative to WT, respectively.

**Figure 4:**
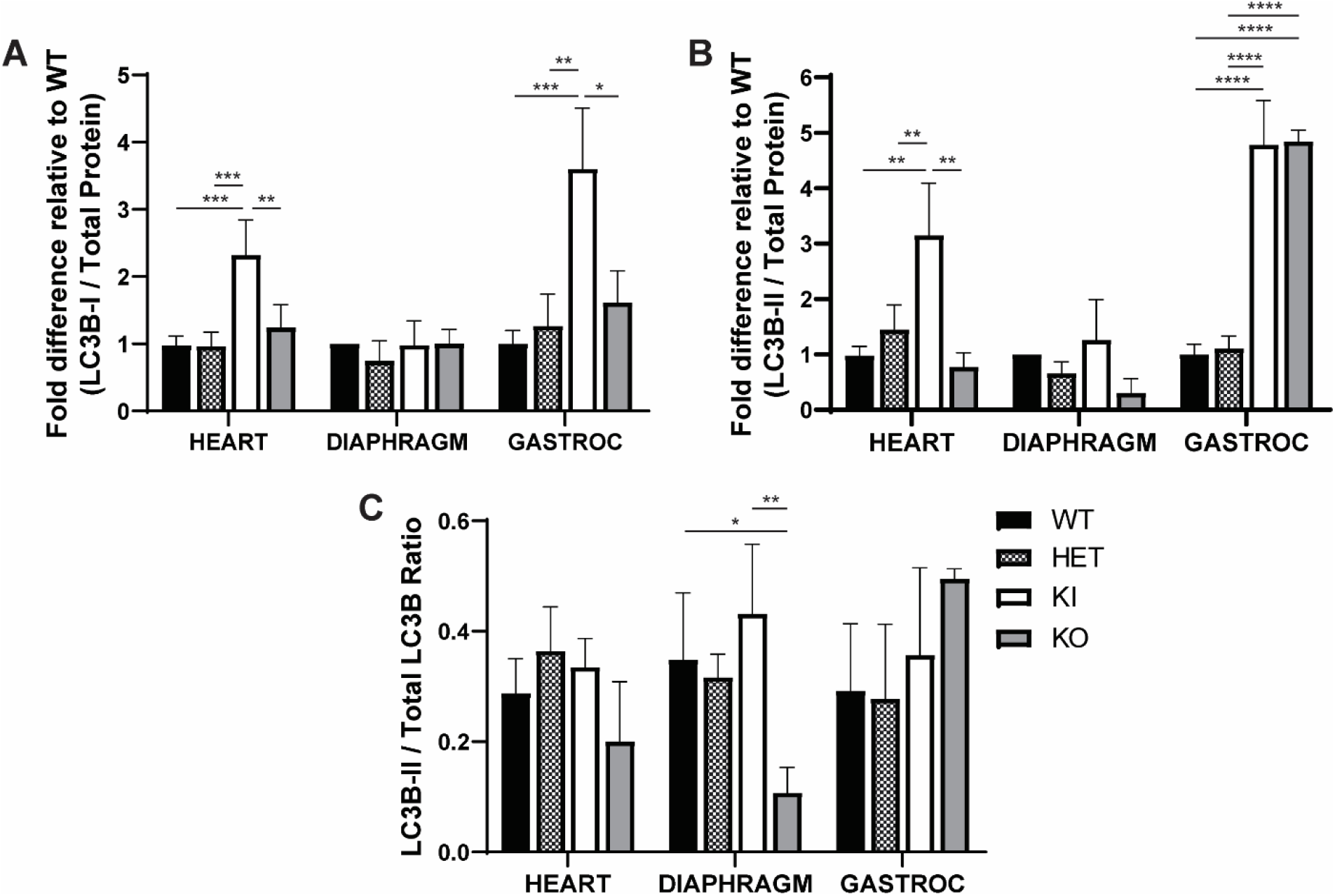
*Gaa*^c.1826dupA^ knock-in mice exhibit markers of impaired autophagy. **(A)** LC3B-I and **(B)** LC3B-II protein expression in the heart, diaphragm and gastrocnemius muscle of *Gaa*^wt^ (WT, n=4), *Gaa*^wt/c.1826dupA^ (HET, n=4), *Gaa*^c.1826dupA^ (KI, n=4) and *Gaa*^tm1Rabn^ (KO, n=3) mice. KI mice display significant increases in LC3B-I protein expression in the heart and gastrocnemius muscle relative to WT, HET and KO mice. KI mice display significant increases in LC3B-II protein expression in the heart relative to WT, HET and KO mice. KI and KO mice both display significant increases in LC3B-II protein expression in gastrocnemius muscle relative to WT and HET mice. Data are reported as fold difference relative to WT expression and are generated from two independent experiments. All comparisons were analyzed using one-way ANOVA with Tukey post-hoc test. ^*^p<0.05, **p<0.01, ***p<0.001, ****p<0.0001. **(C)** LC3B-II to total LC3B ratios in the heart, diaphragm and gastrocnemius muscle of *Gaa*^wt^ (WT, n=4), *Gaa*^wt/c.1826dupA^ (HET, n=4), *Gaa*^c.1826dupA^ (KI, n=4) and *Gaa*^tm1Rabn^ (KO^Rabn^, n=3) mice. Data are reported as absolute ratios of LC3B-II to total LC3B from two independent experiments. All comparisons were analyzed using one-way ANOVA with Tukey post-hoc test. ^*^p<0.05, **p<0.01.

These results suggest that autophagy is impaired in the heart and gastrocnemius muscle, but not the diaphragm, of *Gaa*^c.1826dupA^ knock-in mice by 3 months of age and warrants further monitoring as the disease progresses.

### Cardiac anatomy and function in Gaa^c.1826dupA^ transgenic mice

To assess overall cardiac anatomy and function in *Gaa*^c.1826dupA^ transgenic mice, we performed echocardiography on 3-month old WT, HET and KI mice (**Figure 5A**). Relative to WT and HET mice, KI mice display significant increases in intraventricular septal diameter (IVSd), left ventricular posterior wall diameter (LVPWd) and left ventricle mass index – anatomical hallmarks of hypertrophic cardiomyopathy (**Figure 5B**). KI mice also display alterations in left ventricular internal diameter end systole (decreased; relative to HET) and fractional shortening (increased; relative to WT) – early indicators of abnormal cardiac function (**Supp. Figure 2**).

**Figure 5:**
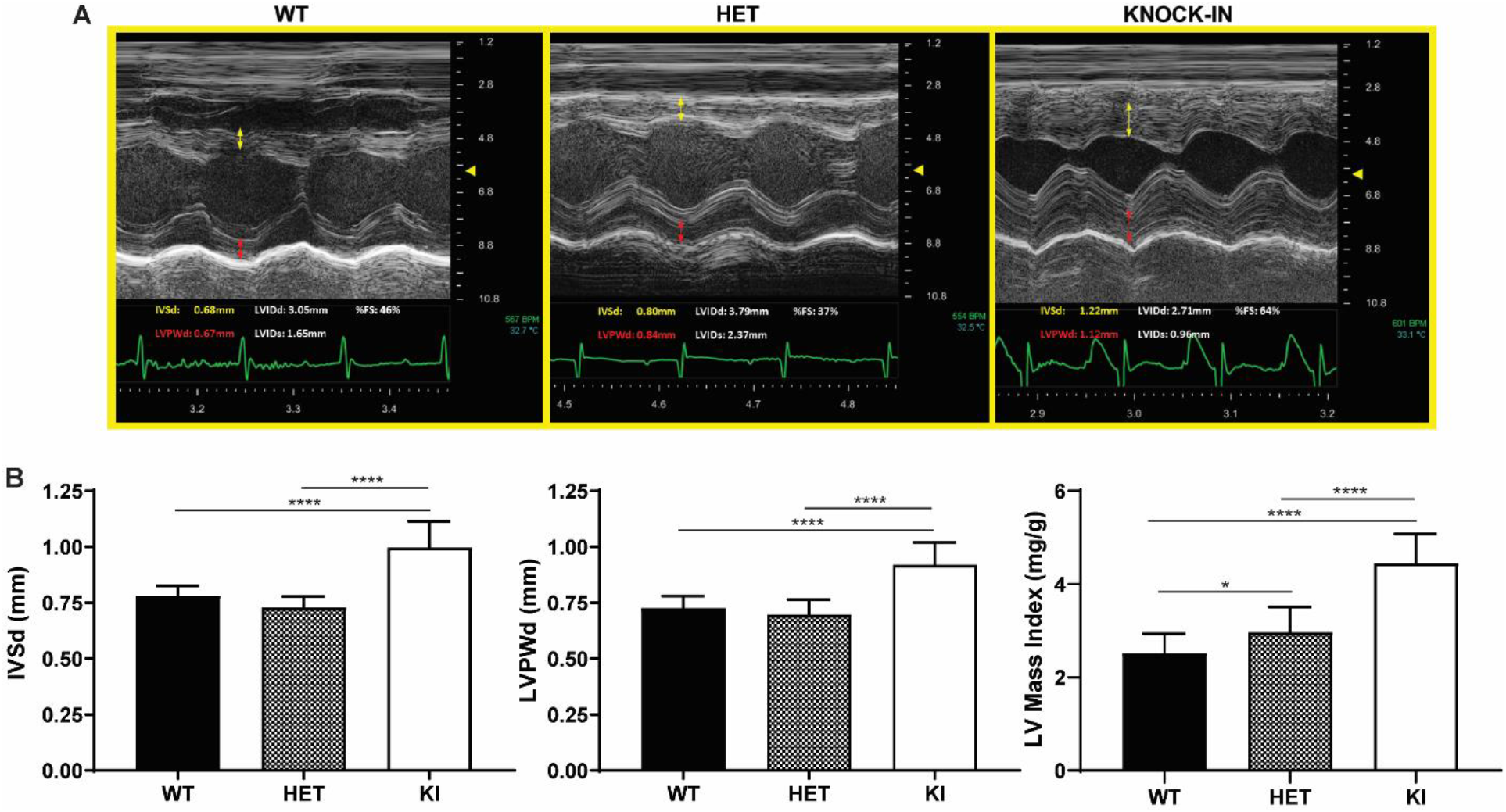
*Gaa*^c.1826dupA^ knock-in mice display anatomical features of left ventricular cardiac hypertrophy. **(A)** Representative echocardiography images of 3-month old *Gaa*^wt^ (WT), *Gaa*^wt/c.1826dupA^ (HET) and *Gaa*^c.1826dupA^ (KI) mice. Interventricular septal diameter (IVSd) is measured in yellow arrows and left ventricular posterior wall diameter (LVPWd) is measured in red arrows. **(B)** IVSd (left panel), LVPWd (middle panel) and LV mass index (right panel) measurements in 3-month old *Gaa*^wt^ (WT, n=27), *Gaa*^wt/c.1826dupA^ (HET, n=12) and *Gaa*^c.1826dupA^ (KI, n=13) mice. Relative to WT and HET mice, KI mice exhibit significant increases in IVSd, LVPWd and LV mass index – anatomical hallmarks of left ventricular cardiac hypertrophy. All comparisons were analyzed using one-way ANOVA with Tukey post-hoc test. *p<0.05, ****p<0.0001.

These results demonstrate that *Gaa*^c.1826dupA^ knock-in mice exhibit early-onset hypertrophic cardiomyopathy – a primary clinical feature of IOPD. Moreover, this work displays the utility of murine echocardiography as a robust diagnostic tool to assess cardiomyopathy in preclinical disease models.

### Forelimb grip strength performance in Gaa^c.1826dupA^ transgenic mice

To assess forelimb muscle strength in *Gaa*^c.1826dupA^ transgenic mice, we measured peak tension force exerted by 3-month old WT, HET and KI mice using a murine-specific grip strength meter. Both male and female KI mice exhibited significant decreases in peak tension force when compared to WT and HET mice as well as a significant reduction in body mass relative to WT mice (**Figure 6**).

**Figure 6:**
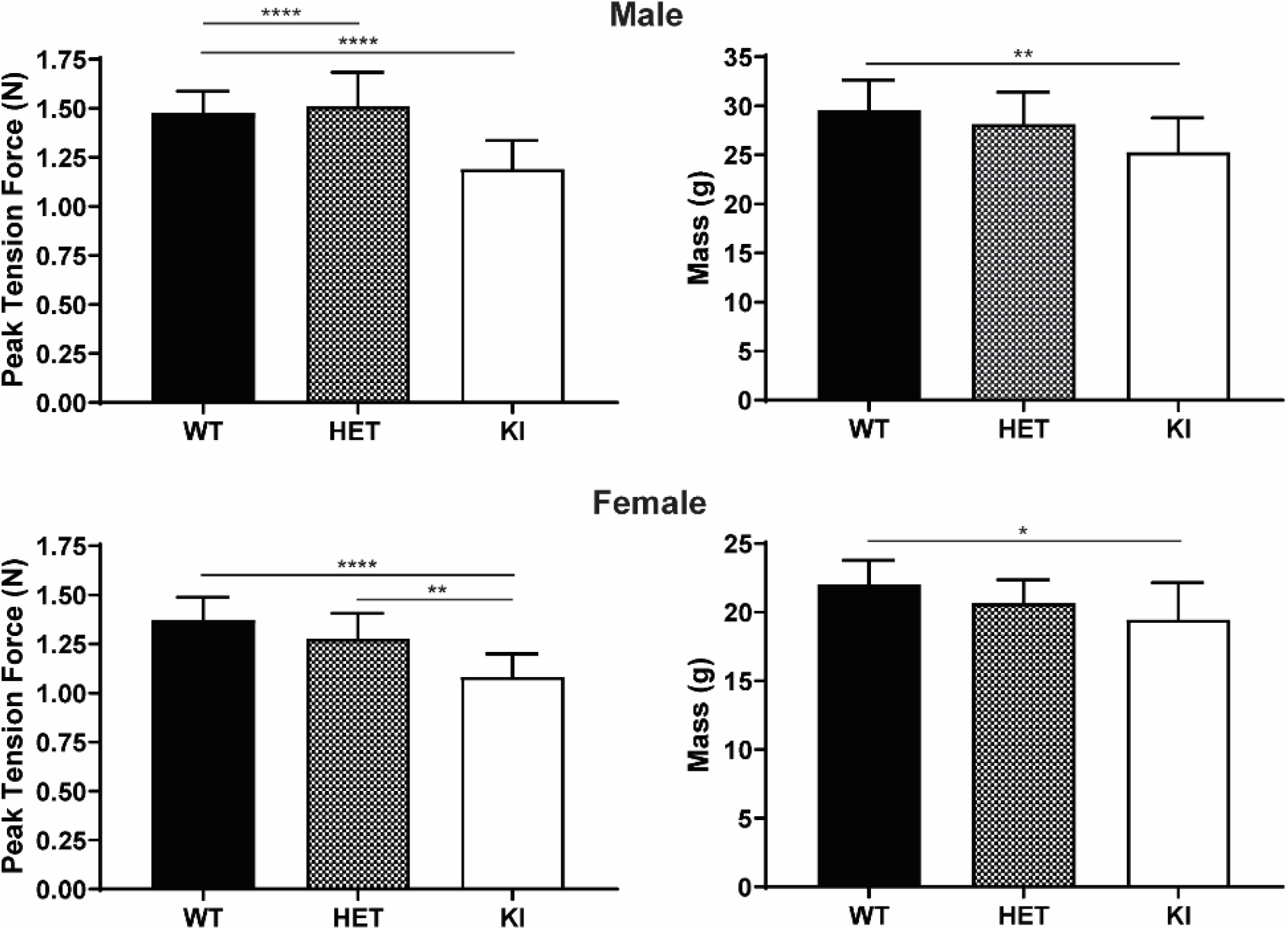
*Gaa*^c.1826dupA^ knock-in mice exhibit decreased forelimb muscle strength. Forelimb peak tension force and body mass measurements in 3-month old male *Gaa*^wt^ (WT, n=12), *Gaa*^wt/c.1826dupA^ (HET, n=10) and *Gaa*^c.1826dupA^ (KI, n=13) mice (**top panels**) and in 3-month old female *Gaa*^wt^ (WT, n=12), *Gaa*^wt/c.1826dupA^ (HET, n=10) and *Gaa*^c.1826dupA^ (KI, n=10) mice (**bottom panels**). KI mice demonstrate significantly reduced forelimb grip strength relative to WT and HET mice and weigh significantly less than WT mice. Forelimb peak tension force was measured using a grip strength meter and taken as average of 9 trials over 3 days. All comparisons were analyzed using one-way ANOVA with Tukey post-hoc test. *p<0.05, **p<0.01, ****p<0.0001.

These results show that *Gaa*^c.1826dupA^ knock-in mice exhibit early-onset musculoskeletal impairment – a key feature of IOPD. *Gaa*^c.1826dupA^ knock-in mice also display a reduction in overall body mass which may be an early indicator of muscular dystrophy.

### Gaa^c.1826dupA^ transgenic mouse histology

To examine cardiac and skeletal muscle tissue structure and glycogen load, we performed periodic acid-Schiff (PAS) staining of heart, diaphragm and gastrocnemius muscle from 3-month old WT and KI mice. In contrast to WT mice, KI mice displayed aggregation of PAS staining in all three tissue types (**Figure 7**). Furthermore, when compared to WT muscle fibers, KI cardiac and skeletal muscle fibers appear irregular in shape and display abnormal extracellular spacing between cells.

**Figure 7:**
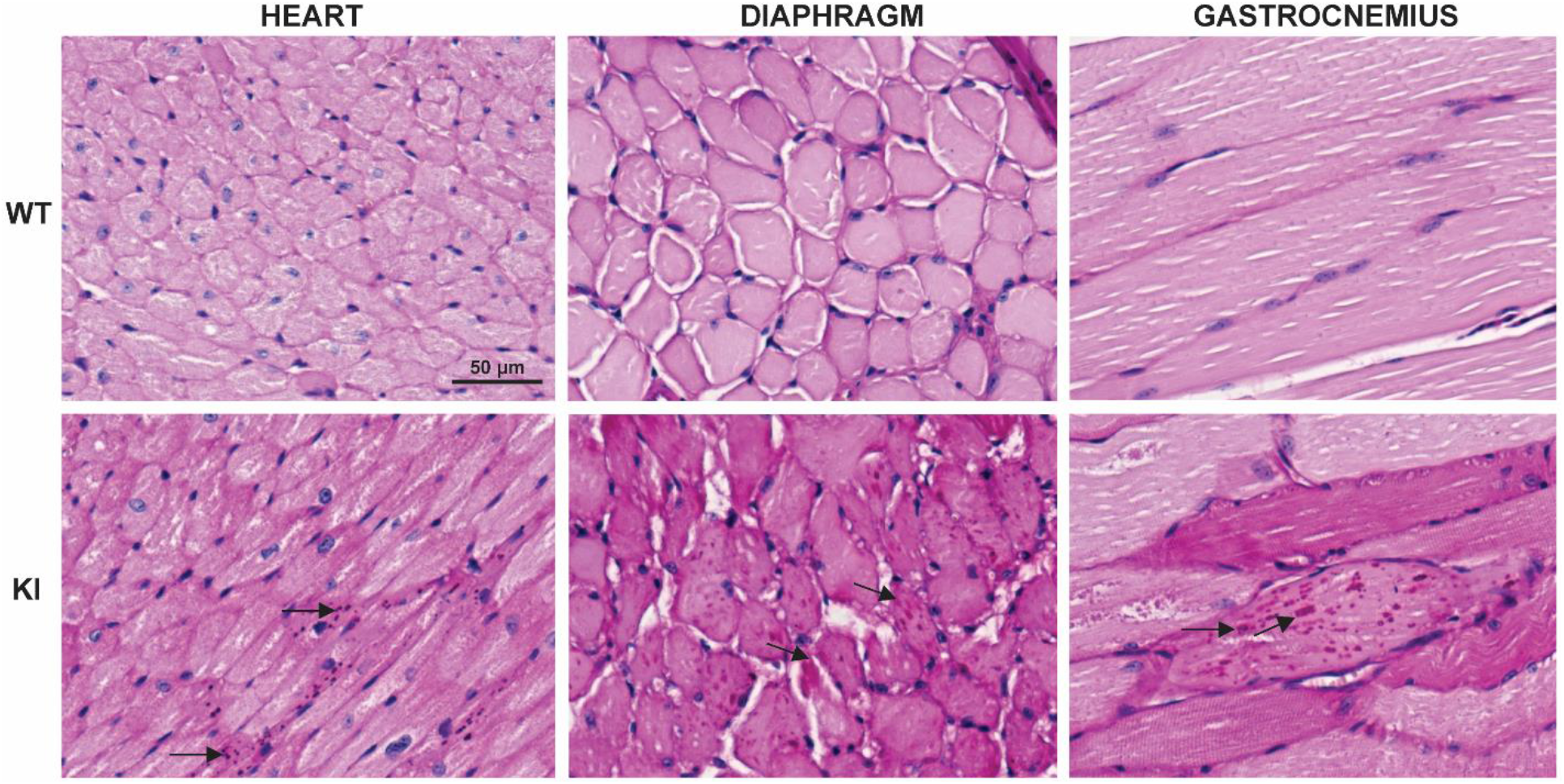
*Gaa*^c.1826dupA^ knock-in mice display abnormal glycogen accumulation in cardiac and skeletal muscle. Representative images of fixed tissue sections from 3-month old *Gaa*^wt^ (WT) and *Gaa*^c.1826dupA^ (KI) stained with periodic acid-Schiff (PAS) stain and hematoxylin counterstain. Following 4% PFA transcardial perfusion, tissue was harvested and either or transversely (heart, diaphragm) or longitudinally (gastrocnemius) sectioned. Images of tissue sections were captured on the EVOS™ M5000 imaging system (Invitrogen) using RGB-mode illumination at 20x objective magnification. Arrows indicate regions of glycogen accumulation.

Altogether, these results show that *Gaa*^c.1826dupA^ knock-in mice display early signs of muscle tissue pathology. The observed PAS-positive aggregates and irregular myocyte structural features are key markers of disease pathogenesis and progression in Pompe tissue.

## Discussion

Currently, preclinical development of novel therapeutic options for Pompe disease rely primarily on the *Gaa*^tm1Rabn/J^ KO mouse model, developed in the late 1990s. While the *Gaa*^tm1Rabn/J^ mouse is an appropriate model for evaluating new enzyme replacement and gene therapy strategies, a Pompe disease KI model bearing a *Gaa* mutation homologous to a known human pathogenic variant is much preferred for development of genome correction-based therapeutics.

This study demonstrates the successful generation of a new KI model of Pompe disease using CRISPR-Cas9 genome editing. Our data show the importance of optimizing HDR-mediated KI efficiency via *in vitro* gRNA and donor template testing prior to *in vivo* application. We found that *in silico* combined rank scoring of gRNAs does not always correlate with actual experimental results. In fact, our *Gaa*^c.1826^ gRNA with the highest predicted rank score (gRNA-1) demonstrated the lowest *in vitro* on-target efficacy. Our data also suggest the enhancement of both gene editing and HDR events by utilizing a dual/multiple gRNA approach versus a single gRNA approach, especially if there is overlap of candidate gRNA target sequences. We found that using a dual gRNA approach, with complete overlap in gRNA target sequences, resulted in the highest level of KI efficiency (12.6%) when compared to each gRNA alone (0 to 7.7%). and was higher than the predicted additive KI efficiencies of the gRNAs (11.2%).

Our preliminary *in vitro* results provide empirical evidence that the dual gRNA approach could increase the probability of generating cellular and murine KI disease models. Consequently, we successfully isolated and characterized a clonal murine C2C12 myoblast line bearing a known pathogenic Pompe disease mutation - *Gaa*^c.1826dupA^ – using the dual gRNA strategy. We found that *Gaa*^c.1826dupA^ KI cells display molecular and biochemical analogy to human Pompe disease with significantly reduced *Gaa* transcript levels, undetectable GAA enzymatic activity and increased glycogen load. Further testing will need to be performed to confirm that this dual overlapping gRNA approach can be broadly applied to increase KI efficiency in other target gene loci and cell lines. *In vivo* application of the dual gRNA approach via pronuclear microinjection of fertilized zygotes resulted in the successful generation of a *Gaa*^c.1826dupA^ KI mouse model which exhibits molecular, biochemical, physiological and histological homology to human Pompe disease. Our *Gaa*^c.1826dupA^ whole genome sequencing results support a recent study^21^ confirming high on-target and undetectable off-target activity in the edited genomes of transgenic mice. Like the *Gaa*^tm1Rabn/J^ KO model, *Gaa*^c.1826dupA^ KI mice display reduced *Gaa* mRNA transcript levels, undetectable GAA enzymatic activity, increased glycogen load and impaired autophagy in multiple tissue types. This disruption in gene expression, enzymatic activity, substrate clearance and autophagy in *Gaa*^c.1826dupA^ KI mice manifests as early-onset cardiac hypertrophy and impaired skeletal muscle function - key phenotypic hallmarks of IOPD. *Gaa*^c.1826dupA^ KI mice also display critical histopathological features of Pompe disease including glycogen accumulation in cardiac and skeletal muscle tissue. Interestingly, *Gaa*^c.1826dupA^ KI mice do not exhibit a lethal phenotype - as observed in untreated IOPD patients - despite development of hypertrophic cardiomyopathy at 3 months of age.

Long-term phenotyping of *Gaa*^c.1826dupA^ KI mice will provide further evidence of analogy to human Pompe disease. We are currently performing longitudinal physiological and histological assessment of *Gaa*^c.1826dupA^ KI mice to determine natural history and disease progression. Given the importance of determining cross-reactive immunogenic material (CRIM) status as it relates to severity of disease progression and immune response to GAA enzyme replacement therapy^22^, we will also aim to determine immune response of *Gaa*^c.1826dupA^ KI mice to recombinant GAA protein.

Altogether, this study provides evidence that *Gaa*^c.1826dupA^ KI mice faithfully recapitulate the IOPD phenotype, with the exception of infantile mortality. While the existing *Gaa*^tm1Rabn/J^ KO mouse model of Pompe disease was also reported to feature reduced GAA expression and enzymatic activity as well as progressive muscle weakness and glycogen accumulation in affected tissue^13^, cardiac hypertrophy – a key distinguishing feature of IOPD – was not fully characterized in *Gaa*^tm1Rabn/J^ mice and was not observed until 32 weeks of age in another knockout model of Pompe disease^12^. Furthermore, the *Gaa*^tm1Rabn/J^ model does not harbor a mutation homologous to one found in human IOPD patients which may ultimately limit its use as a preclinical model for future strategies, especially those targeting the genome. In summary, our results strongly indicate that *Gaa*^c.1826dupA^ mice should be used as the preferred preclinical Pompe disease model for the evaluation of genome correction-based and other genome-targeting therapeutic strategies.

## Materials & Methods

### Gaa^c.1826^ target locus guide RNA and donor ssODN design

*In silico* design of CRISPR-Cas9 guide RNAs (gRNAs) specific for the *Gaa*^c.1826^ target locus was performed using Genetic Perturbation Platform (GPP) sgRNA Designer^23^. Candidate gRNAs were selected using the following criteria: 1) top combined rank score (based upon on-target efficacy and off-target specificity scores) and 2) proximity of predicted Cas9 nuclease cut site to the *Gaa*^c.1826^ target locus. Further potential gRNA off-target analysis was performed using Genome Target Scan (GT-Scan)^19^. Three gRNAs were used in this study (Table 1): gRNA-1 (5’-ACGGCCGGTACGCTGGTCAC-3’), gRNA-2 (5’-GTACGCTGGTCACTGGACAG-3’), gRNA-3 (3’-CTGTCCAGTGACCAGCGTAC-5’).

All donor single stranded oligodeoxynucleotides (ssODNs) were designed with 50bp homology arms flanking the target locus and silent mutations in the protospacer adjacent motif (PAM) and seed region (5nt upstream of PAM) and synthesized by Integrated DNA Technologies.

### Gaa^c1826^ guide RNA spCas9 expression vector cloning

gRNA oligonucleotides with *Bbs*I restriction enzyme overhangs were designed as follows: Forward oligo (5’-CACCG(gRNA)-3’); Reverse oligo (5’-AAAC(reverse complement gRNA)C-3’). Complementary gRNA oligonucleotides were annealed and ligated to *Bbs*I-digested pSpCas9(BB)-2A-Puro plasmid (Addgene plasmid ID# 48139) using Quick Ligation Kit (New England Biolabs). Positive pSpCas9(BB)-gRNA clones were confirmed by Sanger sequencing and expanded using PureLink® HiPure Plasmid Midiprep Kit (Invitrogen).

### In vitro testing of Gaa^c1826^ guide RNAs

pSpCas9(BB)-2A-Puro-*Gaa*^c.1826^ gRNA expression vectors and donor ssODNs were transfected into C2C12 murine myoblast cells (ATCC® CRL-1772™) using nucleofection-based transfection (Neon® transfection system, Invitrogen). For nucleofection-based transfection, 3×10^5^ cells were resuspended in a reaction mixture containing 4.5μg pSpCas9(BB)-2A-Puro-*Gaa*^c.1826^ gRNA expression vector(s) and 450nM ssODN. Cells were electroporated using the following parameters - Pulse voltage: 1650V, Pulse width: 10ms, Pulse number: 3 – and plated onto duplicate wells of Matrigel®-coated 6-well culture plates containing 2mL culture media (DMEM + 10% FBS, 2mM GlutaMax, 100U/mL penicillin, 100μg/mL streptomycin, 0.25μg/mL amphotericin B) and maintained at 37°C with 5%CO_2_. 48h post-transfection, cellular genomic DNA was extracted using QuickExtract DNA solution (Epicentre) and the *Gaa*^c.1826^ target locus was PCR amplified and purified using DNA Clean & Concentrator™ (Zymo Research) and Sanger sequencing was performed (Retrogen). spCas9 nuclease activity and homology-directed repair (HDR) knock-in efficiency were determined by Tracking of Indels by Decomposition (TIDE)^17^ or Tracking of Insertion, Deletions, and Recombination events (TIDER)^18^ analysis of DNA sequence electropherogram files.

### Generation of Gaa^c.1826dupA^ knock-in C2C12 cell line

4.5μg of pSpCas9(BB)-2A-Puro-*Gaa*^c.1826^ gRNA-2 and pSpCas9(BB)-2A-Puro-*Gaa*^c.1826^ gRNA-3 expression vectors (1:1 molar ratio) and 450nM ssODN were added to 3×10^5^ cells and electroporated with similar parameters described above. pCMV6-AC-GFP (OriGene) was used as a transfection positive control and puromycin resistance gene negative control. Nucleofected cells were plated onto duplicate wells of Matrigel®-coated 6-well culture plates for 24 h before adding 2.5μg/mL puromycin dihydrochloride (Sigma-Aldrich) for selection. Puromycin-containing media was replaced every 48h until all pCMV6-AC-GFP transfected cells were no longer viable. Following puromycin-resistant selection, single cell clones were selected by standard serial dilution methods in 96-well plates in the presence of 2.5μg/mL puromycin dihydrochloride. Single cells clones were identified and maintained until sequencing results confirm clonal cell genotype.

### Generation of Gaa^c.1826dupA^ knock-in mice

All study procedures were reviewed and approved under University of California Irvine’s IACUC protocol #AUP 16-63. Pronuclear stage C57BL/6NJ embryos were produced by standard methods^16^. In brief, gRNAs were designed using Genetic Perturbation Platform (GPP) sgRNA Designer and off targets were analyzed by GTScan. 3μM crRNA/tracrRNA/3xNLS-Cas9 protein and 10ng/μL ssODN (IDT) were injected into pronuclei. Surviving embryos were implanted into oviducts of 0.5dpc pseudopregnant ICR females.

### Experimental Animals

The use and care of animals used in this study adhere to the guidelines of the NIH Guide for the Care and Use of Laboratory Animals, which are utilized by the CHOC Children’s Institutional Animal Care and Use Committee. All study procedures were reviewed and approved under CHOC Children’s IACUC protocol #160902.

Heterozygous (*Gaa*^wt/c.1826dupA^) males and females were crossed to obtain homozygous knock-in (KI), heterozygote (HET) and wild type (WT) mice for this study. Experiments were performed on age- and sex-matched mice. Genotyping was confirmed by Sanger sequencing of the *Gaa*^c.1826^ target locus.

### Quantitative real-time PCR

Total RNA was extracted from C2C12 myoblasts or postnatal day 21 tail tip samples using Direct-zol RNA miniprep kit (Zymo Research) and reverse-transcribed using High Capacity cDNA Reverse Transcription kit (Applied Biosystems) following manufacturer’s instructions. TaqMan® Fast Advanced master mix (Applied Biosystems) and specific TaqMan® primer/probe assays for ***Gaa***^c.1826wt^: Forward primer 5’-GGAACACGACCCTTTGTGAT-3’; FAM-hybridized probe 5’ GTACGCTGGTCACTGGACAG-3’; Reverse primer 3’-ATGCAAGATGCTCCCAAGAG-5’ or ****Gapdh**** (TaqMan® assay #Mm99999915_g1) were added to cDNA samples and amplified in triplicate. *Gapdh* was used as an internal reference gene, and relative quantification of *Gaa* gene expression was measured by the comparative ΔC_t_ method.

### GAA enzymatic activity assay

For biochemical analysis, frozen C2C12 myoblast pellets or mouse tissues were homogenized in CelLytic M cell lysis reagent (MilliporeSigma). α-glucosaidase enzyme activity was assessed as previously described with minor modifications^24^. In brief, 10 μL tissue homogenate was mixed with 10 μL of 6 mM 4-methylumbelliferyl-α-D-glucopyranoside substrate (4-MUA, MilliporeSigma) in McIlvaine citrate/phosphate buffer in pH 4.5 and quenched with 180 μL glycine carbonate buffer, pH 10.5 after 1 hr incubation at 37°C in a 96-well plate format. Fluorescence measurements were obtained using an FLx800 spectrofluorophotometer (BioTek) at excitation and emission wavelengths of 360 nm and 460 nm, respectively. One activity unit was defined as 1 nmol converted substrate per hour. Protein concentration was estimated using Pierce BCA assay kit and bovine serum albumin was used as a standard. Specific activity was defined as units of activity per mg of protein.

### Glycogen assay

Tissue glycogen levels were measured using a glycogen assay kit (Sigma-Aldrich) following manufacturer’s instructions. In brief, 10 μL tissue homogenate was incubated with hydrolysis enzyme reaction mixture in a total volume of 50 μL at room temperature for 30 min before adding 50 μL development enzyme reaction mixture for 30 min incubation at room temperature. Absorbance at 570 nm was measured using a spectrophotometer (Multiskan FC Microplate Photometer, Thermo Fisher). A standard curve was generated using standard glycogen solution in the reaction. A reaction without hydrolysis enzyme treatment was used for background correction (endogenous glucose) for each sample.

### LC3B Western blot analysis

Frozen mouse tissues were homogenized in CelLytic M cell lysis reagent (MilliporeSigma) with cOmplete™ protease inhibitors (Roche). Tissue lysates were centrifuged at 10,000 × g for 5 min at 4 °C and supernatants were collected for further testing. Total protein concentration was determined by BCA protein assay (Pierce). 20 μg of total protein lysates in Laemmli sample buffer with 5% β-mercaptoethanol were heated at 95 °C for 5 min. Samples were resolved on 4-15% Mini-PROTEAN® TGX Stain-free™ gels (Bio-Rad) and transferred onto Immun-Blot® PVDF membranes (Bio-Rad). Membrane blots were blocked with EveryBlot blocking buffer (Bio-Rad) for 10 min at room temperature. Blots were then probed with an anti-LC3B primary antibody (rabbit polyclonal, Sigma L7543, 1:1000 dilution) for 2 h at room temperature and a HRP-conjugated goat anti-rabbit secondary antibody (Bio-Rad 1706515; 1:3000 dilution) for 1 h at room temperature. ECL HRP substrate (SuperSignal™ West Pico, ThermoFisher) was added to detect protein targets by chemiluminescence. Stain-free gels and blots were imaged using the stain-free and chemiluminescence settings on the ChemiDoc™ MP imaging system (Bio-Rad). LC3B-I and LC3B-II protein levels were measured by densitometric analysis of Western blots and normalized to the amount of total protein as determined by densitometric analysis of stain-free gels using ImageLab™ software (Bio-Rad). Using the normalized values as determined previously by densitometric analysis, absolute ratios of LC3B-II to total LC3B were calculated as follows: LC3B-II / (LC3B-I+LC3B-II)).

### Periodic acid-Schiff staining

C2C12 cell lines were seeded on Matrigel®-coated 18mm glass coverslips at low density (2.5×10^3^ cells) in culture media at 37°C with 5%CO_2_. 24h post-plating, culture media was replaced with serum-free culture media and maintained for an additional 72 hours with daily replacement of serum-free culture media. 96h post-plating, cells were fixed with 4% paraformaldehyde (4% PFA, Electron Microscopy Sciences) for 30 min at room temperature. Fixed cells were periodic acid-Schiff (PAS) stained (purple-magenta) and hematoxylin counterstained (dark blue) per manufacturer’s protocol (Sigma) and mounted on glass slides with ProLong Gold antifade mountant (ThermoFisher). Representative images were captured on a bright-field microscope (Olympus) at 40x/0.55NA objective magnification.

### Whole genome sequencing and analysis

Whole genome sequencing and analyses were performed on G_0_ wild-type (*Gaa*^wt^), G_0_ founder (*Gaa*^Mosaic^) and G_1_ heterozygous (*Gaa*^wt/c.1826dupA^) tail samples. In brief, 1 μg fragmented genomic DNA was ligated with adaptors using TruSeq DNA libraries and whole genome sequencing was performed on an Illumina HiSeq X Ten Sequencer at 40-50x read depth (Fulgent Genetics). WGS on-target and off-target analysis was analyzed on OnRamp BioInformatics platform. Data were aligned to the Mouse genome (mm10) using BWA^25^. PCR artifacts were identified with the memtest utility from Sentieon^26^, and filtered out using samtools^27^. Alignments were de-duplicated and realigned around insertions and deletions using LocusCollector, Dedup and Realigner from Sentieon. SNV variant calling was performed with GVCFtyper from Sentieon, using the mouse dbSNP 142 data (http://hgdownload.cse.ucsc.edu/goldenpath/mm10/database/snp142.txt.gz) as the known SNPs. Known SNPs and variants falling in un-located chromosomes were removed from the analysis.

For on-target analysis, *Gaa*^c.1826^ target loci from aligned FASTQ reads were designated to the following event categories: 1) *Gaa*^c.1826dupA^ knock-in mutation, 2) insertion/deletion mutation (indel), 3) no mutation, 4) nonspecific mutation. Data are presented as stacked bar graphs indicating percentage of WGS on-target reads for each event category For off-target analysis, we searched for called SNVs that had inserted A, and were flanked by a N->A mutation 4 bases upstream, a N->T mutation 17 bases downstream, and a N->A mutation 20 bases downstream, and then repeated this step for the reverse complement. The only result was the intended *Gaa*^c.1826dupA^ mutation. 5 genomic regions were predicted by GT-Scan to potential be off-target sites of the CRISPR sgRNAs. There were no detected SNVs within 500bp of these sites. The fully processed BAM files (after Realigner) were used as input to the Manta structural variant caller^28^. For each of the non-WT samples, we ran the Manta Somatic caller with the C57BL6-WT sample as ‘normal’ and the sample of interest as ‘tumor’, thereby subtracting the background structural variants in C57BL6-WT compared to mm10. We used vcf-annotate (https://vcftools.github.io) to annotate the output VCF files from Manta.

### Murine echocardiography

Prior to echocardiography, a depilatory cream was applied to the anterior chest wall to remove the hair. 3-month old mice were anesthetized with 5% isoflurane for 15 seconds and then maintained at 0.5% throughout the echocardiography examination. Small needle electrodes for simultaneous electrocardiogram were inserted into one upper and one lower limb. Transthoracic echocardiography (M-mode and 2-dimensional echocardiography) was performed using the FUJIFILM VisualSonics Inc., Vevo 2100 high-resolution ultrasound system with a linear transducer of 32-55MHz. Measurements of chamber dimensions and wall thicknesses were performed. Percentage fractional shortening (%FS) is used as an indicator of left ventricular systolic cardiac function and is calculated as follows: %FS = LVIDd – LVIDs / LVIDd * 100.

### Forelimb grip strength assay

Forelimb grip strength was measured as previously described^29^. One hour prior to grip strength measurement, 3-month old mice were transferred to behavioral room to acclimate subjects to test conditions. Following acclimatization, each mouse was weighed and placed on a forelimb pull bar. Peak tension force exerted by each animal was recorded by a mouse grip strength meter (Columbus Instruments). Each mouse performed 3 pulls per day over 3 consecutive days for a total of 9 pulls per test session. Peak tension force (N) was calculated as the average of each subject’s 9 pulls over the test session.

### Tissue Harvesting, Processing, and Histological Staining

Tail biopsies for genotyping were collected on postnatal day 7 (founder mice) or postnatal day 21 (G_1_ & G_2_ mice). Genomic DNA was extracted using Agencourt® DNAdvance™ genomic DNA isolation kit (Beckman Coulter) with proteinase K and DTT. 3-month old mice were euthanized using CO_2_ asphyxiation and transcardially perfused with phosphate-buffered saline (PBS) followed by 4% PFA for histological staining or PBS alone for biochemical analyses. Heart, diaphragm and gastrocnemius muscle tissue were harvested in this study. Tissue samples for biochemical studies were rapidly frozen and stored at −80°C; tissues for histological staining were processed and embedded in paraffin blocks for sectioning at 4μm thickness and periodic acid-Schiff (PAS) staining was performed as in the above “*Periodic acid-Schiff staining*” Methods section. Representative images were captured on an EVOS™ M5000 imaging system (Invitrogen) using RGB-mode illumination at 20x objective magnification.

### Statistical analysis

All graphs and statistical comparisons were generated using GraphPad Prism 8. Statistical analyses were performed using the two-tailed unpaired t-test or one-way ANOVA followed by Tukey’s HSD test. All data are presented as mean ± standard deviation (SD).

## Funding

This work was supported by the Campbell Foundation of Caring (JYH, SHK, RYW) and the UCLA Intercampus Medical Genetics Training Program Institutional Training Grant Award #2T32GM008243-31 (JYH).

## Author Contributions

JYH and RYW conceived and planned experiments. JYH, SHK, EKS, NDD, ADR, RYW, JN, YC, and JDT performed experimental procedures and critically reviewed the manuscript. JYH and JN designed the Cas9 sgRNAs. JYH designed the ssODN template and *Gaa* spCas9 transfection plasmids. JN performed the murine pronuclear microinjections and subsequent embryo implantations. JYH, RYW, SHK, ND, and JDT wrote the manuscript.

## Competing Interests

JDT is an employee of OnRamp Bioinformatics. The remaining authors have no competing interests to declare.

## Notes

#### Summary of Updates

New figure on LC3B protein expression - a marker of autophagy - in Pompe knock-in model (Figure 4); Author list has been updated to reflect new author (EKS); Figures and Supp. Figures have been rearranged to accomodate new figure

