## Supplemental Tables & Figures for "CRISPR-Cas9 generated Pompe knock-in murine model exhibits early-onset hypertrophic cardiomyopathy and skeletal muscle weakness"

**Table 1:**

| *Gaa*$^{c.1826}$ gRNA | Target Sequence | PAM | Donor Template | On-Target Activity | HDR Efficiency |
|---|---|---|---|---|---|
| 1 | ACGGCCGGTACGCTGGTCAC | TGG | AACACGACCCTTTGTGATCTCCCGCTCAACCTTCTCGGGACATGGCCGGTAA<br>CGCTGGTCACTCGACAGGGGATGTGCGGAGCTCTTGGGAGCATCTTGCAT | 3.8% (±0.1%) | 0% |
| 2 | GTACGCTGGTCACTGGACAG | GGG | AACACGACCCTTTGTGATCTCCCGCTCAACCTTCTCGGGCCACGGACGGTAA<br>CGCTGGTCACTGGACTGGAGATGTGCGGAGCTCTTGGGAGCATCTTGCAT | 22.7% (±3%) | 7.7% (±1.4%) |
| 3 | CTGTCCAGTGACCAGCGTAC | CGG | AACACGACCCTTTGTGATCTCCCGCTCAACCTTCTCGGGCCACGGACGGTAA<br>CGCTGGTCACTGGACTGGAGATGTGCGGAGCTCTTGGGAGCATCTTGCAT | 47.9% (±0.1%) | 3.5% (±2.3%) |
| 2 + 3 | GTACGCTGGTCACTGGACAG<br>CTGTCCAGTGACCAGCGTAC | GGG<br>CGG | AACACGACCCTTTGTGATCTCCCGCTCAACCTTCTCGGGCCACGGACGGTAA<br>CGCTGGTCACTGGACTGGAGATGTGCGGAGCTCTTGGGAGCATCTTGCAT | 41.0% (±4.4%) | 12.6% (±2.9%) |

### Single vs. dual *Gaa*$^{c.1826}$ guide RNA on-target activity and HDR efficiency

Dual overlapping guide RNA strategy (gRNA-2, gRNA-3) achieves the highest HDR efficiency relative to all other guide RNA strategies. Target sequence, PAM motifs and donor templates used for testing of *Gaa*$^{c.1826}$ guide RNAs in C2C12 mouse myoblasts are outlined in table. Desired *Gaa*$^{c.1826dupA}$ knock-in mutation is underlined in red in donor template sequences. All other silent PAM site and gRNA seed region mutations are underlined in donor template sequence in corresponding PAM color. Total on-target Cas9 nuclease activity and HDR efficiency for each *Gaa*$^{c.1826}$ guide RNA condition is displayed as average of two independent experiments.

**Table 2:**

| **Dual gRNA Strategy Components** | **Concentration** | **Founder Mice with any *Gaa* mutation (% Positive)** | **Founder Mice w/ *Gaa*$^{c.1826dupA}$ mutation (% Positive)** |
|---|---|---|---|
| *Gaa*$^{c1826}$ crRNAs (gRNA-2, gRNA-3) | 3uM RNP complex | **66.7%** (8 of 12 founders) | **25%** (3 of 12 founders) |
| tracrRNA | | | |
| 3xNLS spCas9 | | | |
| Donor ssODN | 10ng/uL | | |

**Dual overlapping gRNA strategy and outcomes in *Gaa*$^{c.1826dupA}$ mouse generation**

Dual overlapping guide RNA strategy (gRNA-2, gRNA-3) results in a high percentage of genome-edited founder mice with any *Gaa* mutation (66.7%) as well as the desired *Gaa*$^{c.1826dupA}$ knock-in mutation (25%). Dual overlapping guide RNA components and concentrations used in pronuclear microinjection of C57BL/6NJ fertilized zygotes are outlined in table. Each crRNA was hybridized with tracrRNA at 1:1 ratio to form gRNA duplexes. Equimolar amounts of gRNAs were then combined with 3xNLS spCas9 at 1:1 ratio to form RNP complex. Founder mice positive for any *Gaa* mutation and founder mice with *Gaa*$^{c.1826dupA}$ knock-in mutation are reported as percentages.

**Supp. Figure 1:**

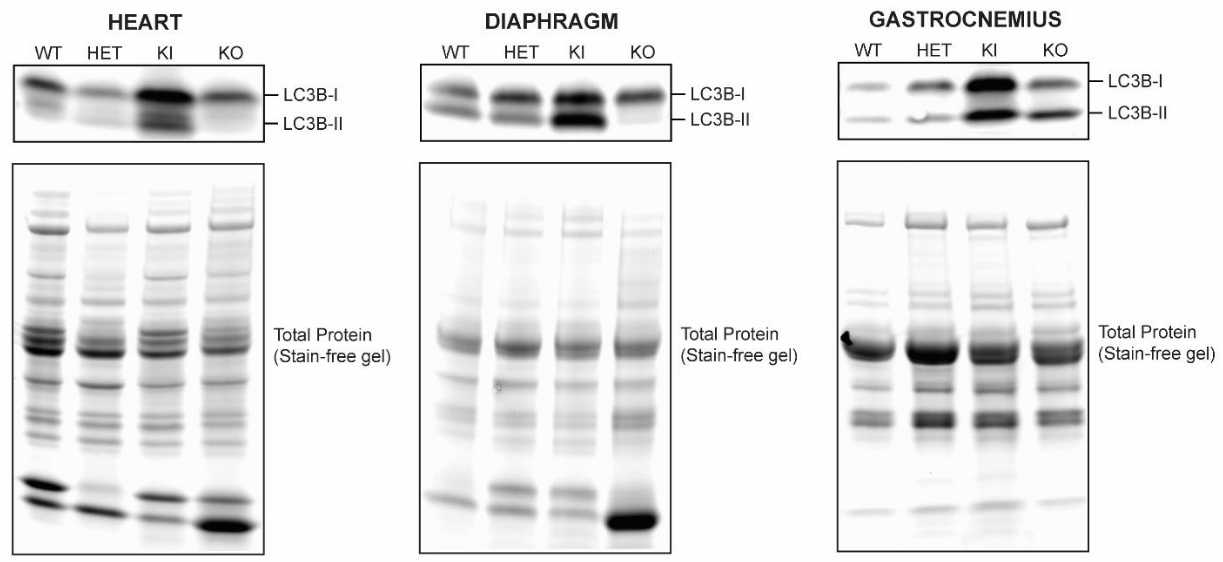


***Gaa*$^{c.1826dupA}$ knock-in mice display increased LC3B-I and LC3B-II protein expression**

Representative anti-LC3B Western blot (top) and stain-free gel (bottom) images of *Gaa*$^{wt}$ (WT), *Gaa*$^{wt/c.1826dupA}$ (HET), *Gaa*$^{c.1826dupA}$ (KI) and *Gaa*$^{tm1Rabn}$ (KO) heart, diaphragm and gastrocnemius tissue protein lysates. LC3B-I and LC3B-II protein levels were measured by densitometric analysis of Western blots probed against a primary LC3B antibody (Sigma, L7543) and normalized to the amount of total protein as determined by densitometric analysis of stain-free gels.

### Supp. Figure 2:

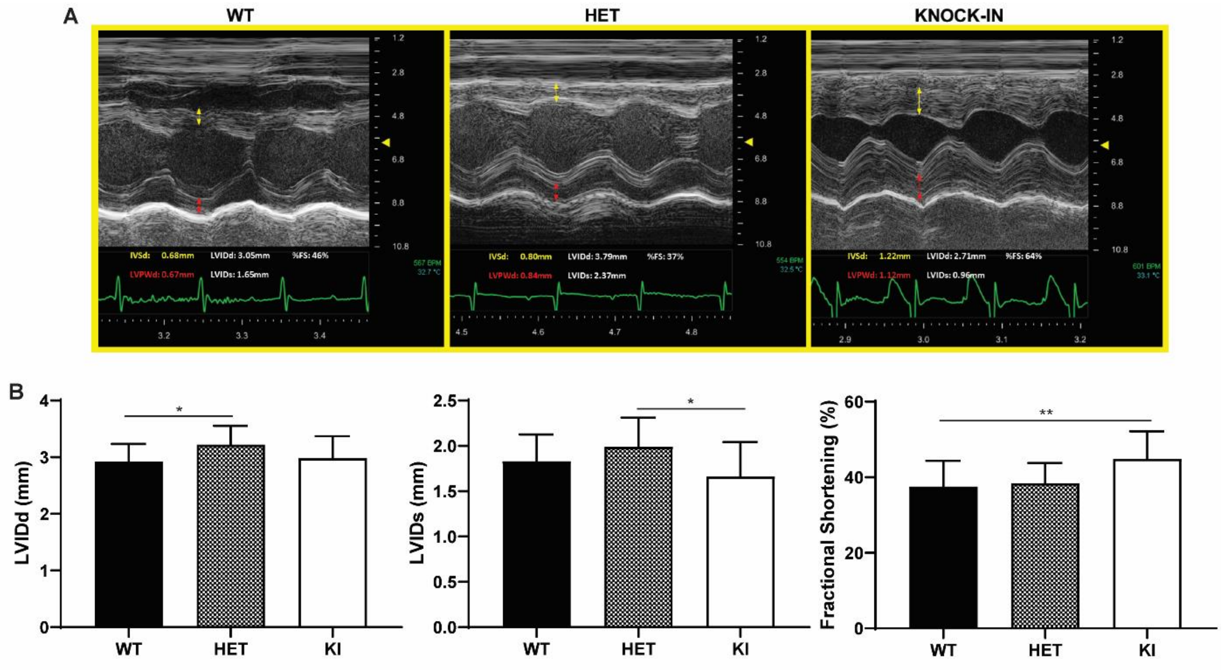


**Gaa$^{c.1826dupA}$ knock-in mice display altered left ventricular cardiac function**

**(A)** Representative echocardiography images of 3-month old *Gaa*$^{wt}$ (WT), *Gaa*$^{wt/c.1826dupA}$ (HET) and *Gaa*$^{c.1826dupA}$ (KI) mice. Additional anatomical measurements include left ventricular internal diameter end diastole (LVIDd), left ventricular internal diameter end systole (LVIDs) and fractional shortening.

**(B)** LVIDd (left panel), LVIDs (middle panel) and fractional shortening (right panel) measurements in 3- month old *Gaa*$^{wt}$ (WT, n=27), *Gaa*$^{wt/c.1826dupA}$ (HET, n=12) and *Gaa*$^{c.1826dupA}$ (KI, n=13) mice. KI mice exhibit significant alterations in LVIDs (relative to HET) and fractional shortening (relative to WT). All comparisons were analyzed using one-way ANOVA with Tukey post-hoc test. *$p<0.05$, **$p<0.01$.
